# Comparing the Effect of Helical-centerline Stent Placement on Blood Flow Velocity with a Straight Stent

**DOI:** 10.1101/2022.05.22.492620

**Authors:** Yutaro Kohata, Hitomi Anzai, Makoto Ohta

**Affiliations:** Graduate School of Biomedical Engineering, Tohoku University, 6-6, Aramaki Aza Aoba, Aoba-ku, 980-8577; The Institute of Fluid Science, Tohoku University, 2-1-1 Katahira Aoba-ku Sendai, 980-8577

**Keywords:** Helical-centreline arterial stent, Angiography images, Blood flow estimation, Time-intensity curve

## Abstract

Stent treatment can be used to treat blood vessel stenosis in a less invasive manner, but re-stenosis is a concern. Because a helical-stent configuration has been thought to reduce the amount of intimal hyperplasia, the helical stent is considered clinically effective. The effects of the helical stent on blood flow velocity, however, have not been studied. In this study, we estimated flow velocities before and after helical stenting using time-intensity-curve (TIC) from angiography images and compared them with straight stenting velocities. As a result, in all cases (N = 3), the velocity reduction was less with helical stenting than with straight stenting. Based on angiography images, this flow estimation method can estimate patient-specific blood flow velocity in situ even in a presence of a stent.

## 1. Introduction

One of the most common treatments for arterial stenosis is a stent. It works by widening narrowed arteries and supplying adequate blood flow to the distal area. A stent can be inserted less invasively with a catheter during an angiography procedure. As a poor prognosis of stent treatment, re-stenosis can occur.

A helical-stent configuration has been proposed, with the stent’s centerline having a helical configuration (1). According to a 2-year follow-up of patients, this helical stent is thought to reduce the amount of intimal hyperplasia (2). Caro et al. identified a helical-stent effect as a swirling of intraluminal flow caused by artery deformation (3). Several studies indicated that vessel wall is affected by wall shear stress which is induced by blood flow (4,5). The flow velocity changes by helical stent then become an appealing issue following helical stent implantation.

Angiography images obtained during the angiography procedure can be used to estimate velocity. The images show the contrast agent flowing along with the patient’s blood flow; the flow velocity can be estimated by analyzing the image contrast changes. The time-dependent change in image intensity is referred to as the time-intensity curve (TIC). Niizawa et al. used TIC from indocyanine green videoangiography to estimate arterial and venous flow velocity in patients (6).

The goal of this study is to estimate flow velocity before and after the placement of a helical stent. To determine the effect of helical-stent placement on velocity, we used TIC to compare helical stenting velocities to straight stenting velocities.

## 2. Method

### 2.1. Clinical setup

Three healthy pigs with both types of stent implants were examined, and an angiography operation was performed as a Case 1, 2, and 3.. A surgical cut down was used to insert a 6Fr femoral access sheath into the right femoral artery. A 0.035-inch guidewire was inserted into both the left and right carotid arteries (LCA and RCA, respectively). A straight stent (ELUVIA 6 mm, Boston Scientific, Marlborough, Massachusetts for cases 1 and 2, and LIFESTENT 6 mm, Bard Peripheral Vascular, Tempe, AZ, the USA for Case 3) and a helical-centerline stent (BM3D 6 mm, Veryan Medical, Horsham, the UK for all cases) were implanted in LCAs/RCAs, respectively.

A biplane angiographic C-arm Infinix Celeve-i 8000C system (Canon Medical Systems, Tochigi, Japan) was used to acquire the images before and after the stent placement. A tip of 5Fr diagnostic catheter was advanced into the LCA/RCA to deliver contrast medium. For each case, several sets of angiography images (1024 × 1024 detecter pixel matrix) were acquired before and after the stent placement (Table 1). Case 3 also had C-arm cone-beam CT image sets acquired before and after the stent placement. All of those medical images were converted into DICOM format.

**Table 1.** Acquired sets of angiography images

| case | number of frames | fps (frame/sec) | resolution (mm/sec) |
| --- | --- | --- | --- |
| 1 | 95 | 15 | 0.174 |
|  | 152 | 15 | 0.174 |
| 2 | 92 | 15 | 0.249 |
|  | 126 | 15 | 0.249 |
|  | 109 | 15 | 0.249 |
|  | 96 | 15 | 0.249 |
|  | 72 | 15 | 0.249 |
| 3 | 177 | 30 | 0.224 |
|  | 187 | 30 | 0.224 |
|  | 189 | 30 | 0.109 |

This Fukushima Medical Device Development Support Center’s Safety Evaluation Department in Fukushima, Japan approved the animal experiments. The protocol numbers are E0000061 for cases 1 and 2 and E0000072 for Case 3.

### 2.2. Velocity estimation

#### 2.2.1. Overview

The following workflow was implemented following the acquisition of angiography images.

#### 2.2.2. TIC method

In this study, the TIC method is divided into two parts: defining regions of interests (ROI) and analyzing TIC. To reduce noise, area-averaged TICs were obtained for each ROI, where the intensity values in each ROI were averaged by the area of each ROI.

First, we extracted vessel areas based on intensity changes in the DSA images. We secondly extracted the centerline of each vessel by applying a thinning algorithm (“sleletonize” function from “morphology” module in python library “scikit-image” (7)) and a smoothing filter (“savgol_filter” function from “signal” module in python library “scipy” (8)) to the extracted vessel areas. We defined two 20 × 20 pixel wide ROIs at proximal and distal points on the centerline.

To reduce noise, image intensity values in an ROI were averaged by the area of the ROI in this study, and an area-averaged TIC was calculated for each ROI. Several local peaks in the area-averaged TICs were confirmed, and the time points for those peaks were automatically detected by finding relative minima/maxima in those TICs (“argrelmax” function from “signal” module in python library “scipy” (8). Figure 1 depicts the area-averaged TICs obtained before and after the helical-stent placement in Case 1.

**Figure 1.**
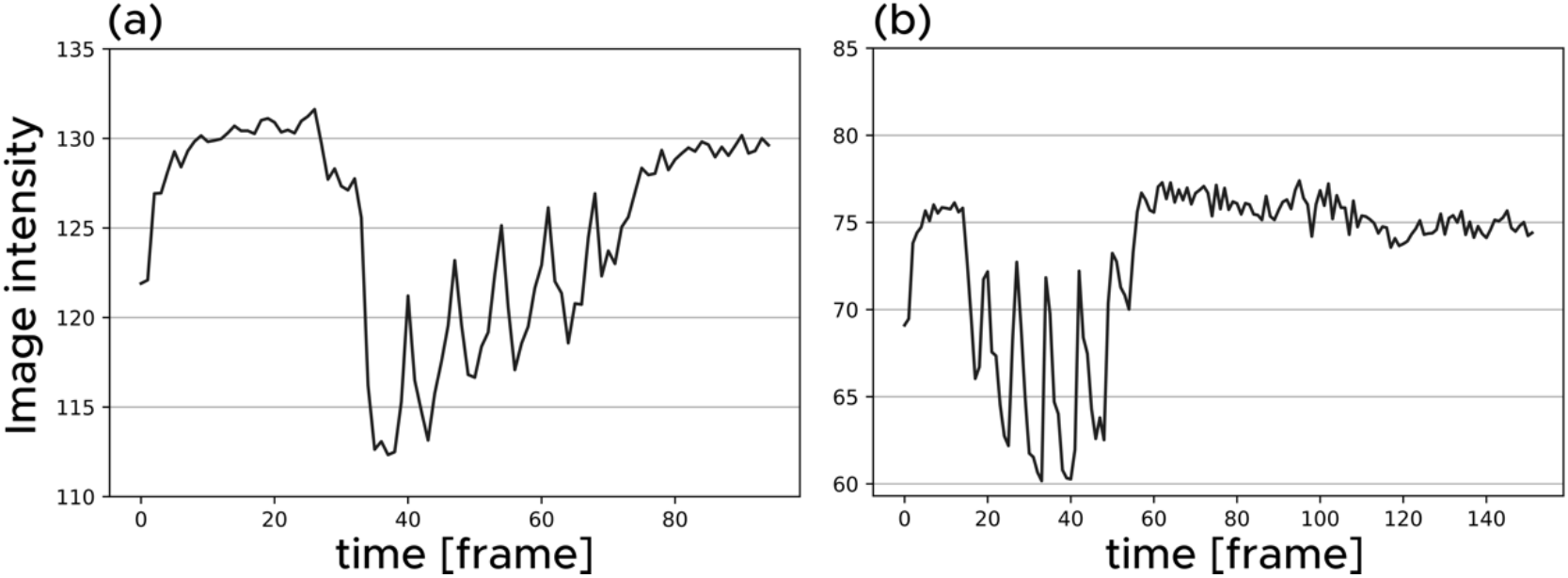
TICs (a) before and (b) after the stent placement for Case 1

#### 2.2.3. Velocity calculation

Along the vessel centreline, the distance between two ROIs, Δx (px), was calculated. For each peak, a time difference, Δt (frame), was calculated by subtracting the peak time at the distal ROI from the one at the proximal ROI. Equation 1 calculated the velocity between two ROIs for each peak in the unit of (mm/s). All peaks’ estimated velocities were averaged, and the averaged velocity was compared before and after stent treatment. The change of the average velocity by stent treatment was evaluated by velocity reduction (%), as calculated in equation 2.

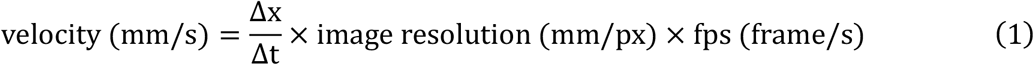

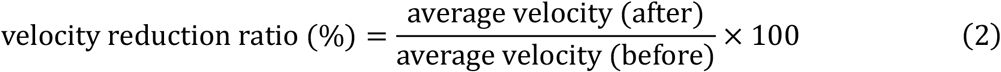

### 2.3. Geometrical analysis

To investigate the geometrical change by stent placements in Case 3, 3D vessel geometries were analyzed using the Vascular Modeling Toolkit (vmtk (9)). First, C-arm CBCT images were used to reconstruct 3D vessel geometry. Second, for each vessel, the vessel centerlines and the radiuses along the centreline were computed.

## 3. Results

The estimated flow velocities for each blood vessel in each case are summarized in Table 2. Flow velocities were reduced in all cases, regardless of stent configurations (straight or helical). The velocity reduction ratio was higher at the helical-stent placement in all cases (N = 3). The reduction ratios were 64.7, 55.0, and 71.3% at helical stenting in each case, while 57.2, 43.0, and 68.0% at straight stenting.

**Table 2.**
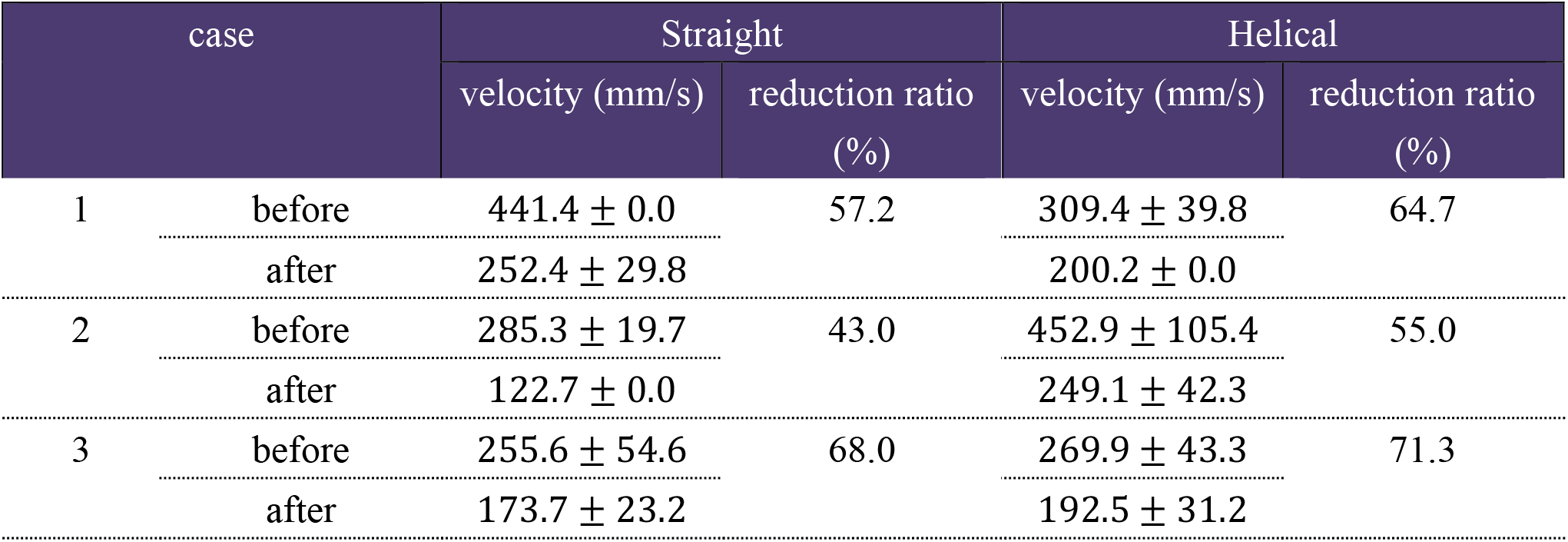
Estimated velocity (mm/s) and velocity reduction ratio (%)

Figure 2 depicts the radiuses along the centerline of each vessel before and after stent treatment in Case 3. The vessel radiuses increased slightly after stent placement, but the increase was less than the resolution of the original image, which is 0.305 mm. In angiography images, a small torsion of the artery was observed after helical stent placement (supplementary material).

**Figure 2.**
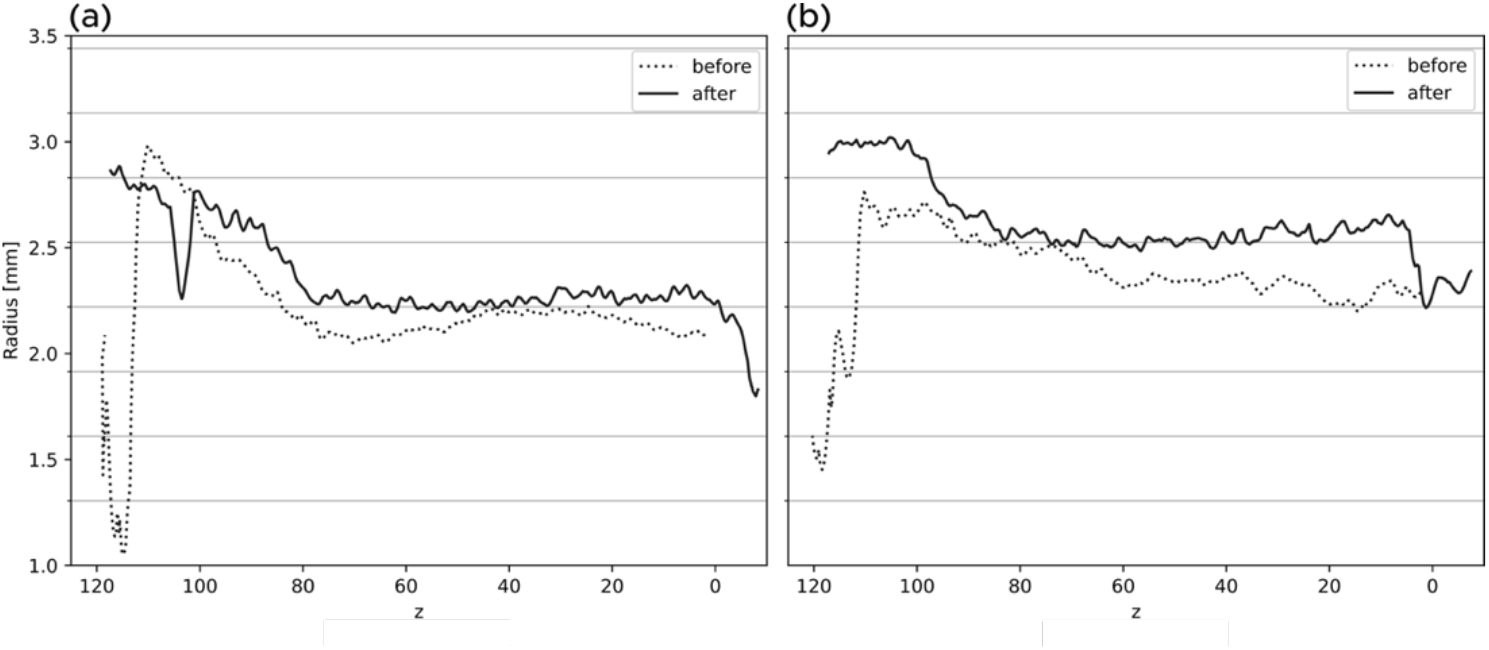
Vessel radiuses along the vessel centerline before (solid line) and after (dotted line) stent teatment in Case 3. (a) straight stent and (b) helical-centerline stent. The mionor y-axis represents the resolution of C-arm CBCT images, which is 0.305 mm.

## 4. Discussion

The flow velocity measurement at helical-stent placement helps to evaluate the helical stent’s swilling flow effect. Using the TIC method, we calculated the flow velocity with stents. In all cases, the reduction ratios of helical stent were less than that of stratight stent. The helical stent may maintain blood flow, and this effect may lead to reduced re-stenosis, as Caro et al. found that the intimal thickness after stent placement was lower in helical stented vessels than in straight stents (3). The reduction ratios at helical stenting and straight stenting were differed between the helical and straight stents. A small torsionby the helical stent may have an effect on the flow velocity.

In all cases, both helical and straight stenting resulted in velocity reductions. Oliveira et al. measured average blood flow velocity in the aorta of pigs and found a reduction in velocity after stent placement (10). This velocity reduction may be related to artery expansion by stent placement(11). However, the expansion in the carotid aretry was less than the image resolution in our study, and determining the effect of stent expansion on vessel radius for both straight and helical stents is difficult.

Using angiography images and the TIC method, we have estimated flow velocities. Angiography images are routinely acquired during the angiography operations. This means that the TIC method can estimate the effect of a stent by comparing patient flow velocity *in situ* just before and after stent replacement. Further research into the frame rate and the ROI distance is still required to validate this method (12). Several peaks in TIC with 15 and 30 fps were observed in this study, and patient flow velocity can be estimated even with a metallic stent.

## Acknowledgements

The authors would like to thank Enago (www.enago.jp) for the English language review.

## References

1. Shinke T, Mullins LP, O’Brien N, Heraty KB, McMahon T, Burke MG, et al. Novel helical stent design elicits swirling blood flow pattern and inhibits neointima formation in porcine carotid arteries. 2008;1.

2. Zeller T, Gaines PA, Ansel GM, Caro CG. Helical Centerline Stent Improves Patency: Two-Year Results From the Randomized Mimics Trial. Circ: Cardiovascular Interventions [Internet]. 2016 Jun [cited 2021 Aug 4];9(6). Available from: https://www.ahajournals.org/doi/10.1161/CIRCINTERVENTIONS.115.002930

3. Caro CG, Seneviratne A, Heraty KB, Monaco C, Burke MG, Krams R, et al. Intimal hyperplasia following implantation of helical-centreline and straight-centreline stents in common carotid arteries in healthy pigs: influence of intraluminal flow ^†^. J R Soc Interface. 2013 Dec 6;10(89):20130578.

4. Malek AM. Hemodynamic Shear Stress and Its Role in Atherosclerosis. JAMA. 1999 Dec 1;282(21):2035.

5. Anzai H, Watanabe T, Han X, Putra NK, Wang Z, Kobayashi H, et al. Endothelial cell distributions and migration under conditions of flow shear stress around a stent wire. Technol Health Care. 2020;28(4):345–54.

6. Niizawa T, Hachiya R, Sugashi T, Terao S, Nagai M, Ishikawa M, et al. Mapping of flow velocity using spatiotemporal changes in time-intensity curves from indocyanine green videoangiography. Microcirculation [Internet]. 2021 May [cited 2022 Apr 6];28(4). Available from: https://onlinelibrary.wiley.com/doi/10.1111/micc.12685

7. van der Walt S, Schönberger JL, Nunez-Iglesias J, Boulogne F, Warner JD, Yager N, et al. scikit-image: image processing in Python. PeerJ. 2014 Jun 19;2:e453.

8. Virtanen P, Gommers R, Oliphant TE, Haberland M, Reddy T, Cournapeau D, et al. SciPy 1.0: fundamental algorithms for scientific computing in Python. Nat Methods. 2020 Mar 2;17(3):261– 72.

9. Antiga L, Piccinelli M, Botti L, Ene-Iordache B, Remuzzi A, Steinman DA. An image-based modeling framework for patient-specific computational hemodynamics. Med Biol Eng Comput. 2008 Nov;46(11):1097.

10. Oliveira JRD, Aquino MDA, Barros S, Pitta GBB, Pereira AH. Alterations of blood flow pattern after triple stent endovascular treatment of saccular abdominal aortic aneurysm: a porcine model. Rev Col Bras Cir. 2016 Jun;43(3):154–9.

11. Mori F, Hanida S, Ohta M, Matsuzawa T. Effect of parent artery expansion by stent placement in cerebral aneurysms. THC. 2014 Apr 1;22(2):209–23.

12. Ruijters D, Benachour N, Grünhagen T, van Nijnatten F, Bonnefous O, Moret J, et al. How to perform DSA-based Cerebral Aneurysm Flow measurements. European Society of Radiology. 2016;20.

