## Supplementary figures and images for "Comparing the Effect of Helical-centerline Stent Placement on Blood Flow Velocity with a Straight Stent"

### supplementary material

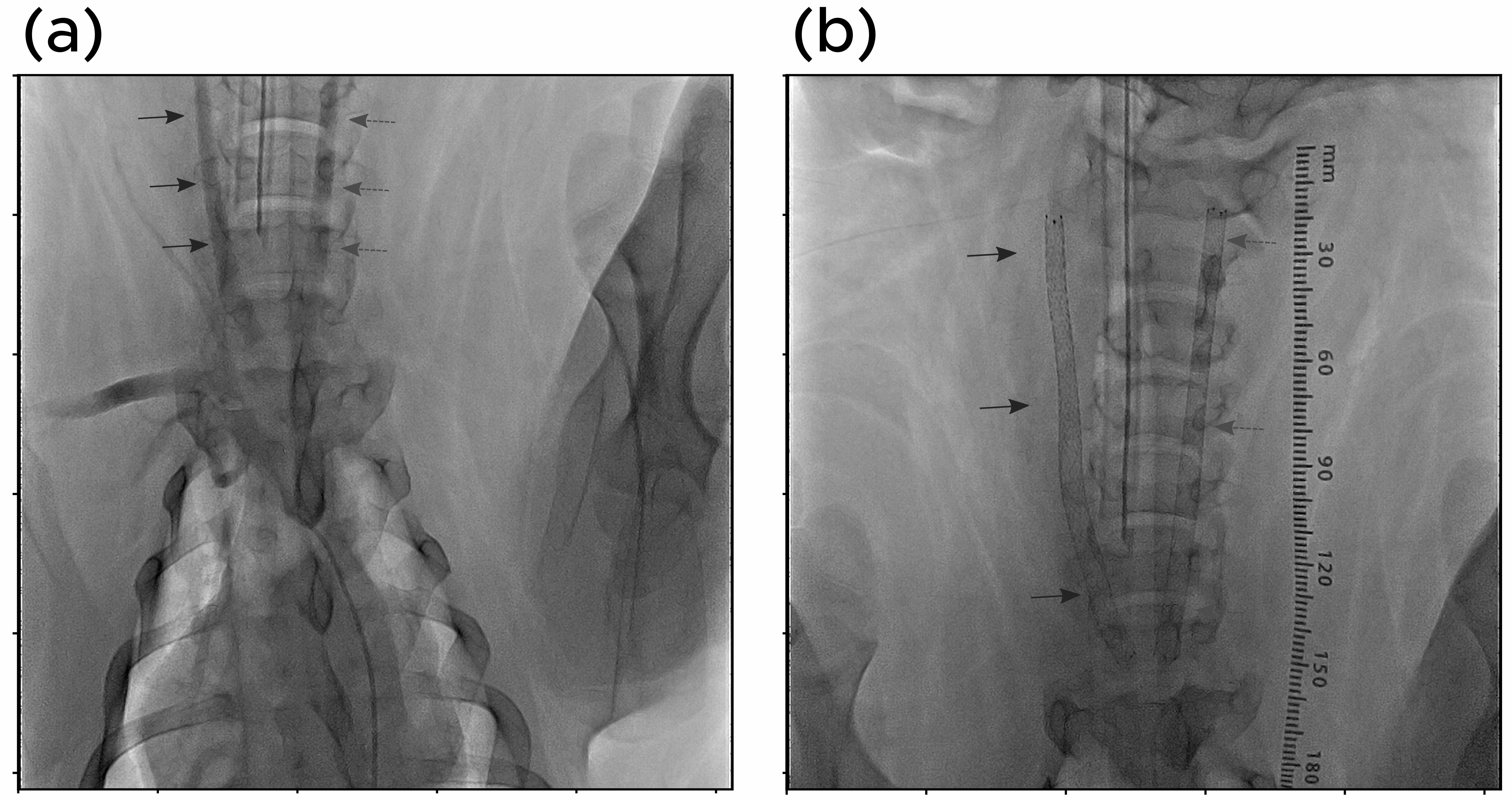
